# Multiscale selection in spatially structured populations

**DOI:** 10.1101/2021.12.21.473617

**Authors:** Hilje M. Doekes, Rutger Hermsen

**Affiliations:** Theoretical Biology, Department of Biology, Utrecht University, Padualaan 8, 3584 CH Utrecht, The Netherlands; Laboratory of Genetics, Department of Plant Sciences, Wageningen University, Droevendaalsesteeg 1, 6708 PB Wageningen, The Netherlands; Centre for Complex Systems Studies, Utrecht University, Leuvenlaan 4, 3584 CE Utrecht, The Netherlands

## Abstract

The spatial structure of natural populations is key to many of their evolutionary processes. Formal theories analysing the interplay between natural selection and spatial structure have mostly focused on populations divided into distinct, non-overlapping groups. Most populations, however, are not structured in this way, but rather (self-)organise into dynamic patterns unfolding at various spatial scales. Here, we present a mathematical framework that quantifies how patterns and processes at different spatial scales contribute to natural selection in such populations. To that end, we define the Local Selection Differential (LSD): a measure of the selection acting on a trait within a given local environment. Based on the LSD, natural selection in a population can be decomposed into two parts: the contribution of local selection, acting *within* local environments, and the contribution of interlocal selection, acting *among* them. Varying the size of the local environments subsequently allows one to measure the contribution of each length scale. To illustrate the use of this new *multiscale* selection framework, we apply it to two simulation models of the evolution of traits known to be affected by spatial population structure: altruism and pathogen transmissibility. In both models, the spatial decomposition of selection reveals that local and interlocal selection can have opposite signs, thus providing a mathematically rigorous underpinning to intuitive explanations of how processes at different spatial scales may compete. It furthermore identifies which length scales—and hence which patterns—are relevant for natural selection. The multiscale selection framework can thus be used to address complex questions on evolution in spatially structured populations.

## Introduction

Spatial structure is the rule, rather than the exception, in biological populations. Examples span a large range of scales. Many bacteria live in biofilms that are highly heterogeneous [1–3] and in which interactions between bacteria are often limited to a range of a few microns [4–6]. At the same time, bushes growing in semi-arid areas form intricate vegetation patterns that span tens to hundreds of meters [7, 8]. Spatial population structure may reflect heterogeneities in the abiotic environment, such as resource availability, but can also arise from self-organisation through ecological interactions between individuals [9].

Because spatial population structure determines with whom organisms interact and compete, it is a key factor shaping evolution. A classical example of this is the evolution of altruism: behaviour that negatively affects an individual’s own fitness, but increases the fitness of its interaction partners [10, 11]. It has long been recognised that a non-arbitrary interaction structure is necessary for altruism to evolve, so that the behaviour of altruistic individuals preferentially benefits other altruists [10, 12–14]. A natural way for such interaction structure to arise is through local interactions and local reproduction, which leads to spatial assortment with altruistic individuals generally being close to other altruists [2, 15–19]. Most formal theoretical work on how spatial structure affects evolution has focused on populations that are divided in distinct groups, *e.g*., trait-groups in which selection is compartmentalised for periods of time [20–22]. Selection is then considered to act at two levels, *within* and *between* groups, and the selection pressures at these two levels can be quantified [20]. Selection pressures within and between groups are not necessarily aligned. In the case of altruism, for example, selection within groups tends to favour selfish behaviour (also called cheating or defecting), while selection between groups promotes altruism [12, 23, 24]. Many biological populations, however, are not subdivided into distinct groups, but are nevertheless structured in space. In such populations, selection can depend on spatial scale, because local selection pressures may differ from those observed at the level of the whole population [25, 26]. Returning to the example of altruism: locally cheaters might out-compete altruists even if altruism is favoured in the population as a whole due to emergent spatial patterning [15, 16, 19]. A formal treatment of selection pressures in such multi*scale* (rather than multi*level*) populations, however, is currently lacking.

Here, we present a mathematically rigorous way to quantify natural selection at different spatial scales. A spatial decomposition of selection is derived which splits global selection into a local component, which describes the average selection *within* local environments, and an interlocal component, which describes the selection *among* these environments. To illustrate the use of this new framework, we apply it to two computational models of the evolution of traits known to be affected by spatial structure: altruism and pathogen transmissibility. We show how the spatial decomposition of selection captures the contribution to selection of processes and patterns at different scales. The multiscale selection framework furthermore allows us to identify which spatial scales are relevant to natural selection, rather than defining those scales *a priori*.

## Results

### A spatial decomposition of selection

Consider a spatially structured population of individuals that differ with respect to some phenotypic value, *ϕ*. This could be a quantitative trait value (*e.g*., an individual’s investment in altruistic behaviour) or an indicator variable that is 1 for individuals that display a certain phenotype (*e.g*., altruism) or possess a certain gene, and 0 for those who do not. We are interested in the evolution of the mean value of *ϕ* over time. Over fifty years ago, George R. Price derived a highly general mathematical description of evolutionary change, showing that the change in mean value of *ϕ* over a given time interval due to selection is equal to the covariance between the phenotypic value and the relative fitness, *w*, of individuals [27]. This covariance, called the *selection differential* and denoted by *S*, provides a general measure of the strength and direction of natural selection.

The selection differential of Price’s equation describes the effect of selection at the level of the whole population. In spatially structured populations, however, this may fail to capture the whole story. For example, consider the hypothetical population in Fig 1a. Here, at the global scale the covariance between phenotype and fitness is positive (black regression line in Fig 1b), yet if the analysis is restricted to individuals within smaller-scale local environments (circles in Fig 1a) it is invariably negative (red lines in Fig 1b). Counter-intuitively, the effect of natural selection can thus be to reduce the mean of *ϕ* in every local environment, while driving it up globally; a spatial Simpson’s paradox [23, 28].

**Fig 1.**
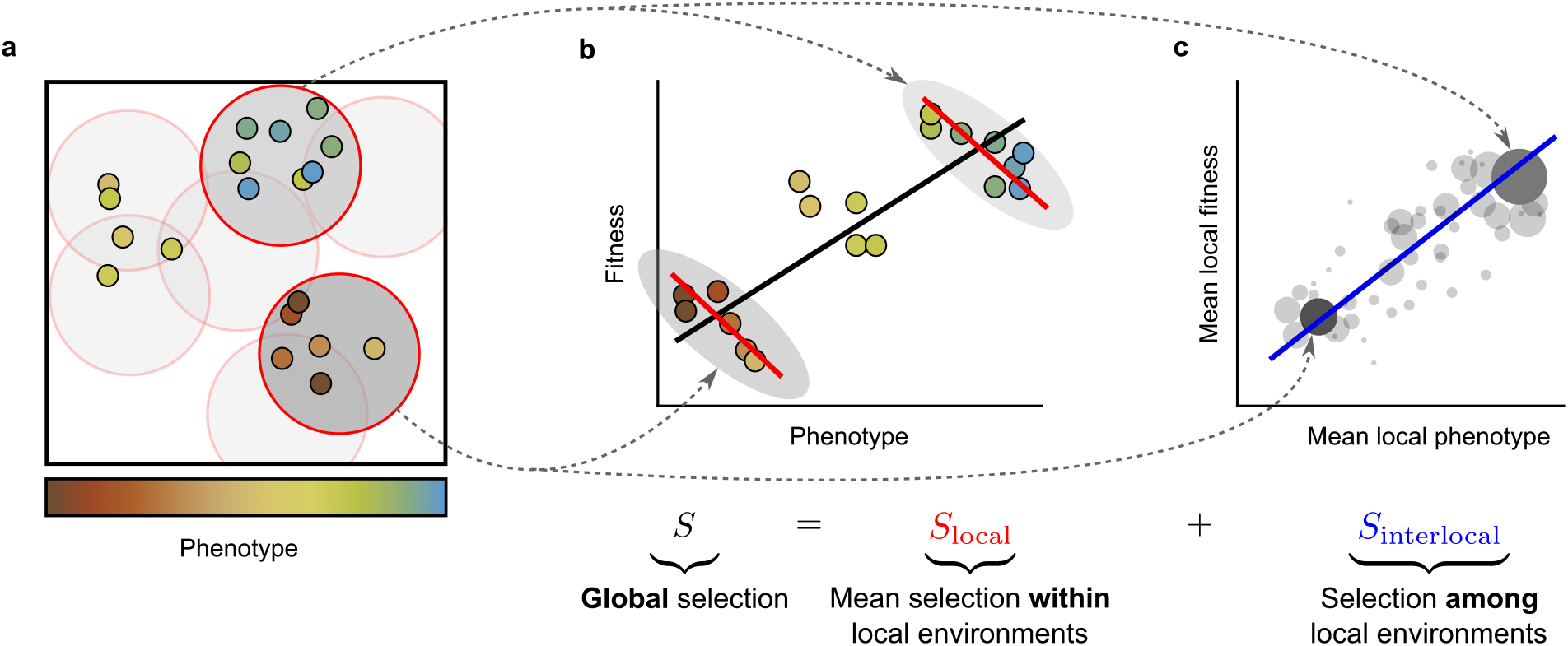
Illustration of the spatial decomposition of selection. (a) Spatially structured population of individuals that differ in some phenotypic characteristic. Local environments are defined as circular areas with a given radius. (b) Example of global and local selection pointing in different directions. The covariance between phenotype and fitness *within* all local environments is negative (*i.e*., local selection is negative), as evident from the negative slopes of the red regression lines; nevertheless, the global covariance between phenotype and fitness is positive (*i.e*., global selection is positive), as apparent from the positive slope of the black regression line. This is an example of Simpson’s paradox. (c) The negative local selection is counteracted by a positive covariance between the mean phenotype and mean fitness of local environments (see the blue regression line). Local environments are weighted by their population density and mean fitness (size of points). This covariance represents the selection *among* environments, *i.e*., the interlocal selection.

To quantify selection at these smaller scales, we first need a mathematically rigorous definition of local environments. In this paper, we simply define local environments as circular areas (disks) with a given radius *r* (see Methods for a more general definition based on a *kernel function*). For any point in space, the local selection at scale *r* can now be measured as the covariance between phenotype and relative fitness of the individuals found within the local environment centred on this point; we call this the Local Selection Differential (LSD; see Methods). Note that local environments may overlap and that they are not necessarily centred on individuals (Fig 1a).

Using this definition of local selection, for any scale *r* we can derive the following spatial decomposition of the selection differential:

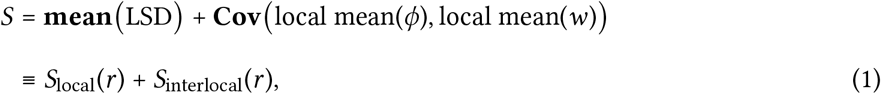

which is the central result of this article (see Fig 1). To properly calculate the mean and the covariance in Eq 1, local environments have to be weighted according to the local density of individuals. A detailed derivation is provided in the Methods; here, we focus on the interpretation of the terms of Eq 1. As the average of the LSDs over all local envrionments, *S*_local_(*r*) describes the *local* component of selection, measuring the selection *within* local environments. The second term *S*_interlocal_(*r*), the (weighted) covariance between local *mean* phenotype and local *mean* fitness, can be interpreted as the *interlocal* component of selection and represents selection *among* environments.

The measure of local selection, *S*_local_(*r*), captures the effect of anything that happens within local environments of size *r*. In other words, it incorporates all mechanisms operating at length scales smaller than or equal to *r*. To identify how a *specific* length scale *r* contributes to selection, we should ask how the local selection component changes if we slightly expand the scale of the local environments from *r* to *r* + d*r*. That is, the contribution to selection of scale *r* is captured by the derivative of *S*_local_(*r*) with respect to *r*, which we denote by *s*(*r*). Assuming that no two individuals can be at the exact same position in space, so that *S*_local_(0) *=* 0, we can then write

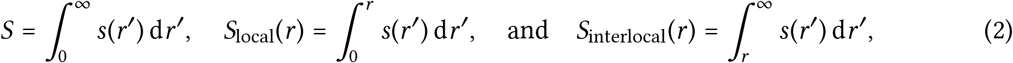

which decomposes *S, S*_local_(*r*) and *S*_interlocal_(*r*) into contributions of different scales.

From Eq 2 we can derive several general insight about the local and interlocal selection components. If local environments become larger and larger, then *S*_local_(*r*) approaches *S*. This makes sense: large “local” environments should capture the global population dynamics. At the same time *S*_interlocal_(*r*) approaches 0, which can be understood by noting that as environments become larger and larger, the local mean values of phenotype and fitness approach the global mean values, eliminating variation between environments. If the environments are made ever smaller (*r* → 0), *S*_local_(*r*) approaches 0 and *S*_interlocal_(*r*) approaches *S*. This also makes sense: very small local environments lack variation in phenotype and fitness, and hence local selection must vanish.

To illustrate the use of the multiscale selection framework (Eq 1–2), we apply it to two models of classical examples of multiscale selection: (i) the evolution of altruism, and (ii) the evolution of pathogen transmissibility.

### Example I: Evolution of altruism aided by self-organising colonies

We model a population of individuals in two-dimensional space that reproduce, die, move around slowly and locally compete for resources (Fig 2a; Supplementary Text). Individuals are characterised by a continuous trait that represents their investment in altruism. Upon reproduction, this trait value is passed on from parent to offspring, at which time mutations are introduced with small probability. Altruistic behaviour directly reduces an individual’s reproduction rate, but benefits all individuals in the local social environment of the altruist. The effects of altruism and resource competition both depend on the distance between organisms, such that an individual that is close to others benefits more from their altruistic action, but also experiences stronger competition (see Supplementary Text for details). The distances beyond which the effects of altruism or resource competition become weak – the “interaction scales” – are denoted by *σ*_a_ and *σ*_rc_.

**Fig 2.**
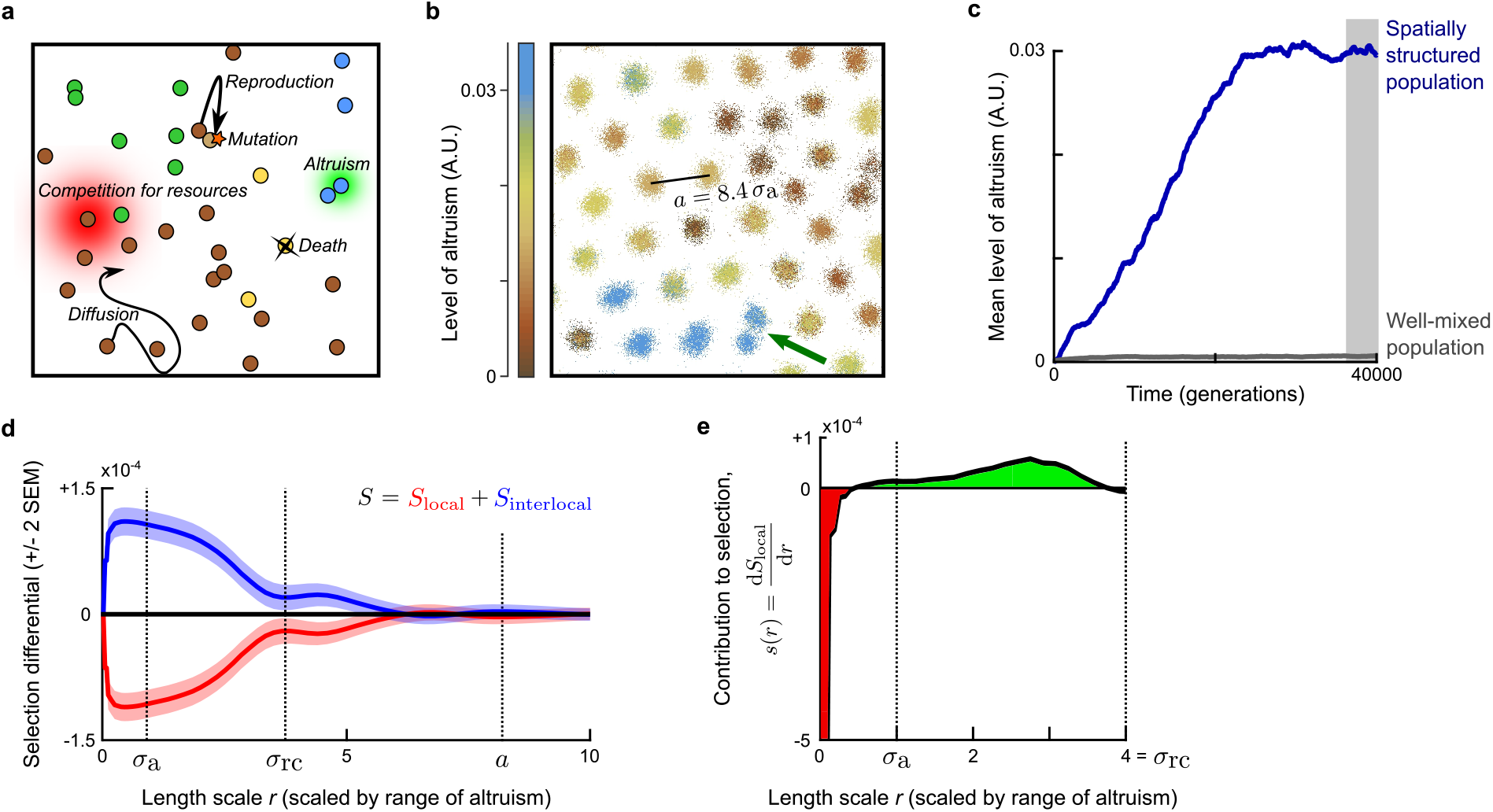
Evolution of altruism. (a) Model illustration. Both resource competition and altruism are local processes. The range of resource competition, *σ*_rc_, is larger than the range of altruism, *σ*_a_. (b) Snapshot of part of the simulation plane (see S1 Movie for dynamics). The hexagonal lattice constant of the emerged colony pattern is *a =* 8.4 *σ*_a_. The green arrow indicates a colony fission event. (c) Mean level of altruism over time, in a population that is well-mixed (grey) or spatially structured (blue). (d) Spatial decomposition of selection differential *S* at evolutionary equilibrium for varying length scales of the local environments. Space was scaled such that the range of altruism *σ*_a_ = 1. *S*_local_(*r*) and *S*_interlocal_(*r*) were calculated as averages over 10 000 instances of the simulation plane obtained between Time = 36 000 and 40 000 generations (shaded area in panel c). (e) Contribution to selection of different length scales. Red areas indicate a negative contribution to selection, green a positive contribution.

If the interaction scale of altruism, *σ*_a_, is sufficiently smaller than the scale of resource competition, *σ*_rc_, the model population shows intriguing self-organisation (Fig 2b, S1 Movie): a Turing-like instability results in a hexagonal pattern of distinct colonies that display Darwinian dynamics of their own. In colonies with a high mean level of altruism, the density of individuals is high because they all benefit from the altruism of colony members. Over time, however, the level of altruism within a colony declines because mutants with lower levels of altruism are selected (“defectors” or “cheaters”). This decline eventually results in the demise of the colony, after which it is replaced by a newly-formed colony that originates from the binary fission of one of the surrounding colonies (see arrow in Fig 2b, S1 Movie). Crucially, colonies with a higher mean level of altruism are more likely to reproduce. These emergent colony dynamics are studied in depth in a companion paper [29].

The simulations are initialised with individuals that do not invest in altruism. But, over the course of the simulation, the mean level of altruism increases until it reaches a stable value (Fig 2c). In contrast, if we destroy the self-organised pattern by mixing the population (*i.e*., randomly assigning positions to individuals every time step), altruism does not evolve at all (Fig 2c). The emergent spatial patterns are hence crucial for the evolution of altruism, consistent with previous modelling work [15–17, 30, 31].

Once the mean level of altruism has stabilised, we should expect the global selection differential to equal zero on average because no directional selection remains. To average out fluctuations arising from the stochastic dynamics in the finite population, we take the mean of the selection differential over 4 000 generations (shaded area in Fig 2c), and indeed find that over this period, *S* ≈ 0 (black line in Fig 2d). However, the spatial decomposition of selection of Eq 1 reveals a completely different picture (Fig 2d). For *r*-values up to six times the altruism interaction range, the mean local selection, as measured by *S*_local_(*r*), is negative. This negative local selection is compensated by positive interlocal selection, *S*_interlocal_(*r*). These results capture and formalise the verbal explanation given above: within local environments, individuals with a lower level of altruism are selected because they benefit from the altruistic behaviour of others nearby while paying less costs; but local environments in which the mean level of altruism is low also tend to have low mean fitness, *i.e*., selection *among* local environments favours higher levels of altruism.

Fig 2d illustrate how *S*_local_(*r*) and *S*_interlocal_(*r*) depend on the radius of the local environments, *r*. The effects observed for small and large *r*-values represent general properties of the spatial decomposition of selection (see Eq 2): for large *r, S*_local_(*r*) converges to the global selection differential (which is close to zero in this case) and *S*_interlocal_(*r*) converges to zero, while for small *r, S*_local_(*r*) declines because the variation in phenotype and fitness within the local environments is reduced.

Intuitively, one might expect that the scale associated with negative selection on altruism is tied to the size of single colonies. The shortest distance between colonies is given by the lattice constant of the emerging hexagonal colony pattern, *a =* 8.4*σ*_a_ (see Fig 2b). Indeed, for local environments with radius 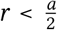 (such that most disks contain individuals of one colony only), *S*_local_(*r*) is clearly negative (Fig 2d). However, when we consider the contribution of each length scale to selection, *s*(*r*), we see that only very small length scales of *r* < *σ*_a_ contribute negatively to selection, while length scales of *r* ⪆ *σ*_a_ contribute positively (Fig 2e). This indicates that colonies are not homogeneous: even within a single colony we observe assortment of individuals with different investment in altruistic behaviour, and this assortment contributes positively to the selection of altruism. This is understandable: individuals that are very close together (distance *< σ*_a_) experience a similar level of altruism and competition. In very small local environments (*r < σ*_a_), cheaters hence must have an advantage over altruists because they pay less costs. But once the scale of local environments become similar to the range of altruistic interactions, individuals within the same local environment may experience different levels of altruism, and the effects of these heterogeneities start to contribute to local selection.

Looking further into which spatial scales are relevant for selection, we see that the scale for which the differences between *S*_local_ and *S*_interlocal_ vanish is close to the lattice constant *a* (Fig 2d). This indicates that local environments that are large enough to contain individuals from more than one colony capture the processes and patterns contributing significantly to global selection are; larger-scale patterns, such as the clear assortment at the colony level (see Fig. 2b), appear to have a negligible effect.

In conclusion, the multiscale selection framework allows us to mathematically show that local selection for altruism is negative, but that this is compensated by positive interlocal selection. It furthermore provides a way to quantify how specific length scales contribute to selection, thus revealing which patterns and processes are significant for natural selection. Specifically, it shows that the heterogeneity of colonies is relevant to selection, whereas the assortment at the colony level is not.

### Example II: Evolution of pathogen transmissibility in an SI-model

As a second example, we consider the evolution of the transmission rate of an endemic pathogen in a spatially structured population of host individuals. This model is rooted in a long tradition of such epidemiological models; see *e.g*., [32–36].

In the model, host individuals live on a 2D square simulation lattice (Fig 3). They can be either susceptible to infection, or infected. Susceptible individuals reproduce asexually, in which case the offspring is placed on a neighbouring lattice site. Each lattice site can hold at most one individual; susceptible individuals therefore locally compete for empty space. Infected individuals do not reproduce, and they die at a higher rate than susceptible individuals. The pathogen is transmitted locally at a rate that varies among pathogen variants. We consider the evolution of this pathogen transmissibility. For simplicity, each infected individual is considered to carry a single pathogen strain, and mutations instantaneously change the transmissibility of all pathogens within a single infected host (*i.e*,, newly arising pathogen variants rapidly sweep the within-host pathogen population). Details on the model implementation are provided in the Supplementary Text.

**Fig 3.**
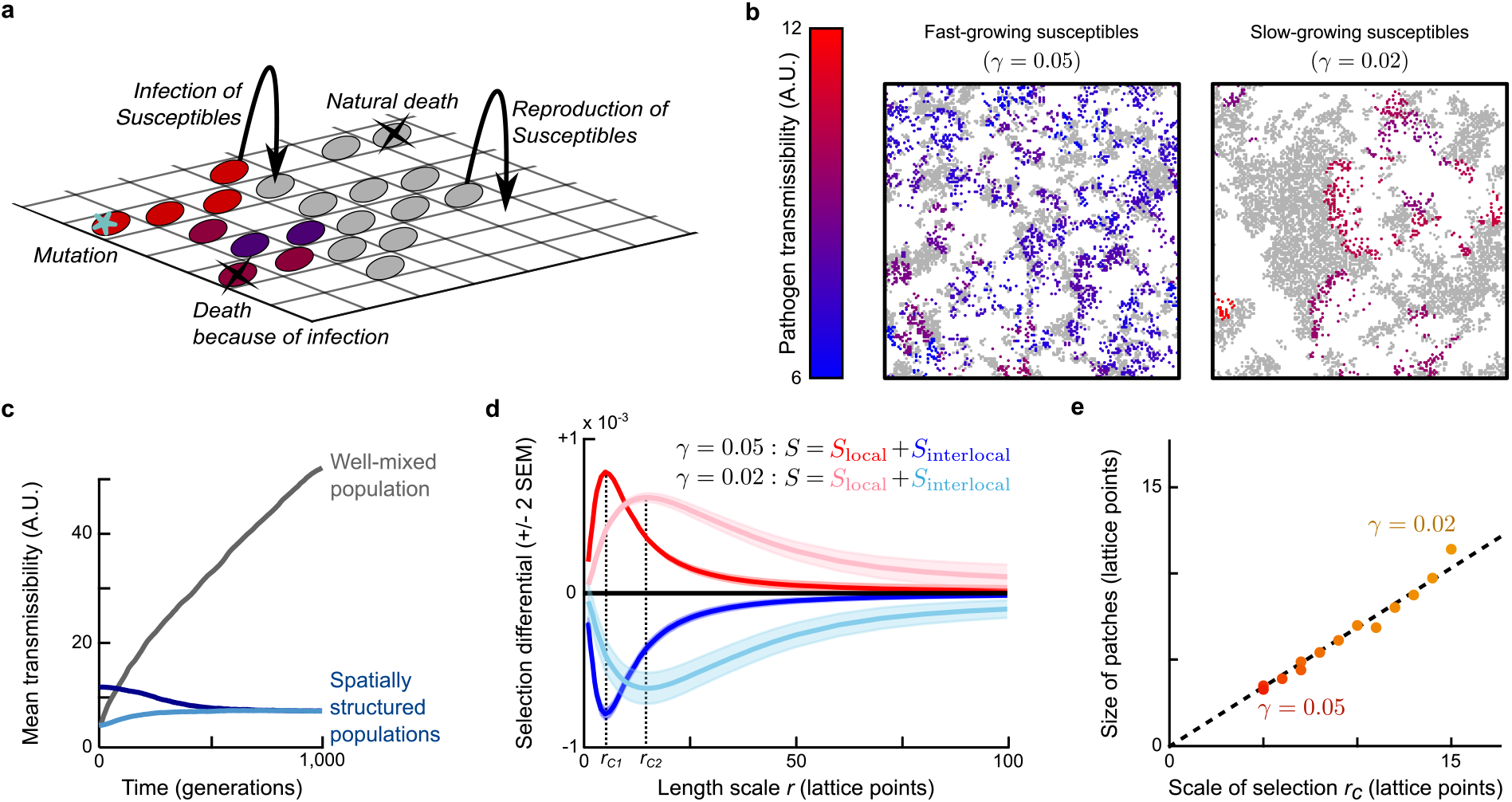
Evolution of pathogen transmissibility. (a) Model illustration. (b) Snapshot of part of the simulation lattice for two different values of the reproduction rate of susceptible individuals, *γ*. Susceptible individuals are plotted in grey, infected individuals are coloured based on the transmissibility of the pathogen they carry. See S2 Movie and S3 Movie for dynamics. (c) Mean transmissibility of the pathogen over time in populations that are well-mixed (grey) or spatially structured, with different initial transmissibility values (blue) (d) Spatial decomposition of selection differential S at evolutionary equilibrium for varying length scales of the local environments. For both values of *γ, S*_local_ and *S*_interlocal_ were calculated as averages over 10 000 instances of the simulation lattice obtained between Time = 9 500 and 10 000 generations. We define the critical scale of selection, *r*_C_, as the length scale at which the contribution to selection switches from positive to negative (*i.e*., where *s*(*r*) *=* d*S*_local_*/*d*r* switches sign). (e) Critical scale of selection, *r*_C_, plotted against size of the emerged patterns for different values of the susceptible reproduction rate *γ*. Pattern size was determined using the pairwise correlation function (see Supplementary Text).

After initialisation, the simulated population quickly self-organises into spatial patterns: the infection chases patches of susceptible individuals in wave-like structures (Fig 3b, S2 Movie and S3 Movie). These patterns strongly influence the evolution of pathogen transmissibility (Fig 3c): If pattern formation is prevented by constantly mixing the population, pathogens with ever-increasing transmissibility are selected because pathogens with higher transmissibility spread faster among the available susceptible hosts. In spatially structured populations, however, the mean transmissibility eventually stabilises (blue lines in Fig 3c). This is explained by a feedback between evolution and the emergent spatial patterns: pathogen strains shape their local environment, and this environment in turn affects the strain’s local fitness [9, 25, 37]. Specifically, pathogens with very high transmissibility are affected by ‘self-shading’ [34, 38]: they rapidly deplete the susceptible hosts in their vicinity and are then left with little opportunity to spread, locally resulting in low average pathogen fitness. In contrast, more prudent pathogens shape their environment in such a way that sufficient susceptible hosts remain available to allow the infection to continue spreading [35, 39].

We use the multiscale selection framework to quantify and formalise this self-shading. As before, we allow the population to reach evolutionary equilibrium and then calculate *S*_local_(*r*) and *S*_interlocal_(*r*) over a range of *r* values (Fig 3d). While the global selection differential *S* is insignificant as expected (because the global mean transmissibility no longer changes and mutations are unbiased), within local environments selection still favours pathogens with high transmissibility: *S*_local_(*r*) > 0 for small *r* (red line in Fig 3d). This effect is counteracted by negative interlocal selection: *S*_interlocal_(*r*) < 0 for small *r* (blue line). The negative interlocal selection confirms that pathogens with higher transmissibility more often reside in local environments in which susceptible host availability and hence pathogen spread is limited. We can now further explore how the spatial patterns affect the evolutionary process. The size of the spatial patterns that emerge in the population depends on several model parameters [35] including the reproduction rate of the susceptible hosts: lower reproduction rates result in larger patterns (Fig 3b). To demonstrate how these larger patterns are reflected in *S*_local_ and *S*_interlocal_, we repeated our analysis with a lower susceptible reproduction rate (right panel in Fig 3b, pink and light-blue lines in Fig 3d). Evidently, the curves representing *S*_local_(*r*) and *S*_interlocal_(*r*) are stretched towards larger scales. This reflects that the spatial scales relevant to selection depend on the size of the patterns in the population: after all, selection for pathogen restraint can only be observed if the local environments in which selection is measured are large enough to cover multiple patches.

To explore this relationship between multiscale selection and spatial pattern size, we define the critical scale of selection *r*_*C*_ as the length scale at which the contribution of length scales to selection, *s*(*r*), switches sign (Fig 3d), such that scales smaller than *r*_*C*_ contribute positively to selection, and scales larger than *r*_*C*_ contribute negatively. This critical scale of selection is an emergent property of the dynamics. By repeating the analysis for a range of susceptible growth rates, we find that the critical scale of selection is proportional to the size of the emergent patterns (Fig 3e). Hence, the *S*_local_- and *S*_interlocal_−curves, and specifically the critical scale of selection *r*_C_, capture the length scale of the spatial structures that are relevant for natural selection.

In conclusion, this second example illustrates that the multiscale selection framework can be used to measure and quantify self-shading. It furthermore allows the identification of the scale(s) of population structures relevant to natural selection.

## Discussion

We have presented a new, multiscale selection framework that can be used to analyse evolution in spatially structured populations. The framework is based on a spatial decomposition of selection (Eq 1–2) that quantifies local and interlocal selection for any spatial scale. Two example models illustrated how this framework can be used to measure the contribution to selection of processes and patterns at varying scales, and thus to identify the spatial scales relevant to natural selection.

The spatial decomposition of selection demonstrates how natural selection in local environments can substantially differ from the selection in the global population, even if an average is taken over all local environments. Evolutionary studies based on local observations can hence provide an incomplete picture of the evolutionary dynamics in the global population. The framework presented here provides a way of determining whether a collection of local sampling areas is representative of the whole population under selection. Namely, if the local sampling areas are large enough to capture all spatial structures and processes relevant to selection, interlocal selection vanishes and the mean local selection differential (*i.e*., as measured within sampling areas) is equal to the global selection differential. Specifically, this means that there should be no covariance between mean phenotype and mean fitness among local environments (*i.e*., *S*_interlocal_ *=* 0). A significant correlation between local mean phenotype and local mean fitness is a strong indication that larger structures exist in the population that contribute to natural selection. However, to reliably estimate *S*_local_ and *S*_interlocal_, observations from many individuals are required, especially if selection is weak. While such data is easily obtained in simulation studies, it remains to be seen if the framework can also be successfully applied to observational or experimental data.

The examples of multiscale selection studied here—the evolution of altruism and of pathogen transmissibility— were chosen because they are among the best-known examples of a feedback between spatial patterns and eco-evolutionary dynamics, and because this feedback has also been confirmed experimentally. For instance, increased population viscosity facilitates the evolution of altruistic public good production in lab populations of Pseudomonas aeruginosa [40, 41], and several experiments have shown that increased host population viscosity and/or localised pathogen spread select for lower virulence in an insect larval virus [38] and bacterial viruses [42, 43]. The effect of spatial structure on natural selection is, however, by no means limited to these two examples. It was first described in models of catalytic hypercycles of self-replicating molecules, which give rise to self-organised rotating spirals that select for higher death rates of the individuals constituting these spirals [44, 45]. Since then, many other examples have been described [9, 25], including anticompetitor toxin production in bacteria [46–50]. Applying the multiscale selection framework to such other examples could lead to new insights in these systems as well.

The spatial decomposition of selection in Eq 1 is analogous to the decomposition of selection into within- and between-group components for distinct, non-overlapping groups derived by Price in 1972 [20]. It however provides a different perspective on the effects of (spatial) structure on selection. The multilevel framework (*i.e*., the distinct-groups approach) requires that distinct and non-overlapping groups are defined. If the spatial structures are variable in time and space, such as the patches in the model of pathogen transmissibility, this is infeasible and the multiscale approach is more appropriate. If distinct groups can be easily recognised, such as in the altruism model, the group selection framework can provide important insights into the selection pressures acting at different levels of organisation. This is explored in depth in [29]. But even in this case the multiscale approach may provide additional insights, for instance by showing that relevant structure exists within groups or in the spatial organisation of the groups themselves.

While the multiscale decomposition of the selection differential is new and uniquely untangles local and interlocal selection in spatially structured populations, it is not the only, nor the only correct way the selection differential can be decomposed. Next to the distinct-groups multilevel approach discussed above, the selection differential can also be decomposed into terms that capture the effect of an individual’s own character on its fitness, and the effect of its environment (the contextual analysis approach to multilevel selection) [51], or in terms that capture the effect of an individuals behaviour on its own fitness and on the fitness of related interaction partners (the inclusive fitness framework) [26, 52, 53]. Each decomposition of the selection differential tells a potentially new story about the underlying mechanisms driving evolution. The multiscale selection framework presented here is an addition to the toolbox available to address complex evolutionary questions.

## Methods

Below, we derive Eq 1. Full specifications of the simulation models, as well as computational details on the calculation of the two terms of Eq 1, are provided in the Supplementary Text.

### Background: Definition of the selection differential *S*

Consider a population in space that at some time *t* consists of *n* individuals. Let *ϕ*_*i*_ be the phenotype of individual *i*, and *W*_*i*_ its fitness, defined as the number of offspring at some later time *t* + Δ*t*, including the individual itself if it survived over the time step Δ*t*. Price’s equation [27] now states that

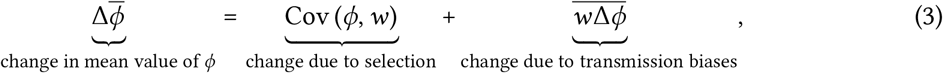

where 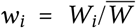 is the relative fitness of individual *i* over the time interval (we use the common notation 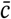 to denote the population mean of characteristic *c*) and Δ*ϕ* captures biases in the transmission of phenotypic values from parents to offspring. Importantly, the first term of the Price’s equation reveals that the effect of selection on the mean phenotype is captured by the covariance between phenotype and fitness; this term is also called the *selection differential*. The analysis presented here focuses on the selection differential only; for a full version of the Price equation with overlapping generations, including transmission and survival-bias effects, see *e.g*., [54].

### Measuring selection in a local environment: the Local Selection Differential (LSD)

For any point ***m*** in space, the local population density is defined as a conventional kernel density estimate:

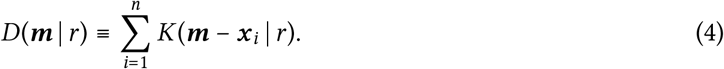

Here ***x***_*i*_ is the position of individual *i*. Its contribution to the density at position ***m*** depends on its distance to position ***m*** according to the kernel function *K*(***y***|*r*). The parameter *r* is the scale parameter (or band width) or the kernel function. In effect, the kernel function defines the local environments: it determines which organisms contribute to what extent to the environment around each point in space.

In this paper, we use disk-shaped kernel functions which include an individual in the local environment only if its distance to the environment’s midpoint is smaller than a given radius, *r*. Other reasonable choices for the kernel function include bi-variate normal or exponential distributions, which weigh individuals close to the focal point more heavily than those further away.

A proper kernel function is normalised (*i.e*., the integral of *K*(***y*** | *r*) over space is equal to one); it follows that

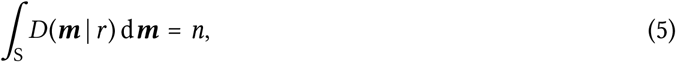

where *∫*_S_ represents the integral over the entire space.

For any characteristic *c* of individuals, such as phenotype or fitness, we define the *local mean* at point ***m*** as

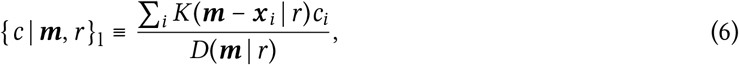

which is the average of *c* over all individuals, weighted by the kernel function based on their position relative to ***m***. We will often write {*c}*_l_ as a shorthand for {*c* | ***m***, *r}*_l_ to avoid clutter.

Analogous to the selection differential in Price’s equation, we can now define the local selection around point ***m*** as the covariance between phenotype and relative fitness within the local environment, which we call the *Local Selection Differential* (*LSD*):

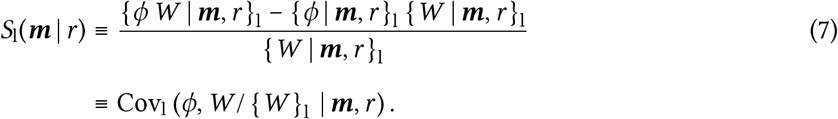

Note that the LSD is equal to the local covariance between *ϕ* and local relative fitness (*W /* {*W* }_l_), *i.e*., the fitness of an individual relative to others in the local environment. Thus defined, it measures the effect of selection on the change in the local mean of *ϕ*at position ***m***, {*ϕ*| ***m***, *r}*_l_.

### Decomposing the selection differential into local and interlocal selection

For any function over space *g*(***m***), we define the *spatial mean* as

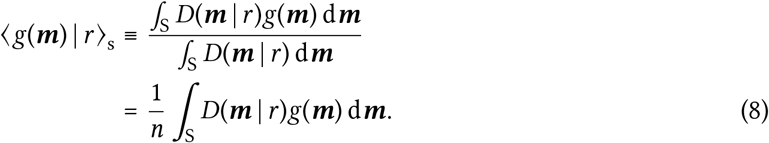

Note that this represents the average of *g*(***m***) over the complete space, but that each position ***m*** is weighted by the local density *D*(***m*** | *r*). This weighting is equivalent to the weighting by group size in Price’s derivation of within- and between-group selection in a population consisting of distinct groups [20, 21]. For readability, we will often write ⟨*g*(***m***) ⟩_s_ for ⟨*g*(***m***) | *r*⟩_s_. Conveniently, given the definitions of Eq 6 and 8, the spatial mean of the local mean is simply the population mean, *i.e*.,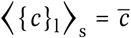.

Using these definitions, can now derive the desired decomposition:

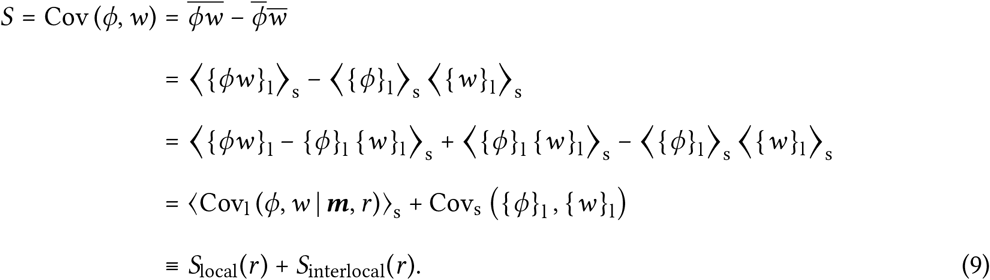

Hence, *S*_local_ is the average of the LSD over all local environments (defined by their midpoints ***m*** and scale *r*), where each environment is weighted by its local density and its local mean fitness:

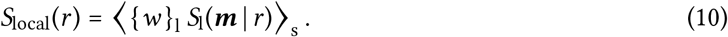

It captures the total effect of selection *within* local environments. On the other hand, *S*_interlocal_(*r*) is the spatial covariance (defined in terms of spatial means) of the local mean phenotype and local mean fitness. This term captures the selection *among* environments.

## Code availability

Simulation codes and analysis scripts are available from GitHub: https://github.com/rutgerhermsen/altruism.git (altruism model) and https://github.com/hiljedoekes/MultiscaleSelection_SI (SI model).

## Acknowledgements

We thank Reinder Bosman for a preliminary analysis of the SI-model, and Laura van Schijndel for valuable discussions and comments on the manuscript. This work was supported by the Human Frontier Science Program, grant nr. RGY0072/2015 (http://www.hfsp.org/funding/research-grants).

## Author Contributions

R.H. conceived the project with input from H.M.D. The mathematical derivations were performed collaboratively by both authors. R.H. developed and analysed the altruism model with input from H.M.D.; H.M.D developed and analysed the SI-model with input from R.H. H.M.D. wrote the first draft of the manuscript, which was then substantially edited by both authors.

## Supplementary Information

### Supplementary Movies

**S1 Movie. Dynamics of the altruism model**. Simulation lattice points are coloured by the mean level of altruism of the individuals at that position, according to the colour scale shown in the movie and in Fig 2b. Empty lattice points are white. A high-quality version of this video is shared here: https://doi.org/10.5281/zenodo.5608897.

**S2 Movie. Dynamics of the SI-model if reproduction of susceptibles is fast (***γ =* 0.05). Empty lattice points are coloured black, susceptible individuals white, and infected individuals are coloured according to the transmissibility of the pathogen they carry (see colour scale in Fig 3b).

**S3 Movie. Dynamics of the SI-model if reproduction of susceptibles is slow (***γ* = 0.02). Same colour scheme as S2 Movie.

### Supplementary Text

#### Individual-based model of the evolution of altruism

As a first example of multiscale selection, we consider the evolution of altruism in a spatially explicit individual-based model. A full analysis of this model is provided in a companion paper [29].

#### Model description

We model a population of individuals living in a 2D space with periodic boundary conditions. Each individual can reproduce asexually, die, and move in an undirected fashion. Individuals differ by one continuous phenotype, *ϕ*, that specifies their level of altruism. By itself, altruistic behaviour is costly: it directly reduces the individual’s rate of reproduction. However, an altruistic individual does improve the living conditions of all individuals in its local neighbourhood (including itself), irrespective of their phenotype. Individuals locally compete for resources: the rate of reproduction of an individual decreases with the density of individuals in its local neighbourhood.

Both altruistic interactions and competition are modelled through a bi-variate normal (Gaussian) interaction kernel function *G*(***y*** | *σ*), where *σ* is the standard deviation of the kernel. In terms of this function, we denote the total level of altruism experienced by individuals at position ***y*** by

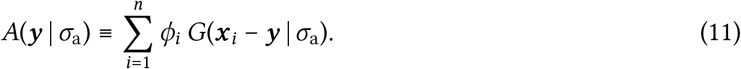

Here, *n* is the population size. Although we do not intend to model a particular altruistic behaviour, it is convenient to envision *A*(***y*** | *σ*_a_) as the availability of a public good that is locally deposited by altruistic organisms. The scale *σ*_a_ can be interpreted as the interaction range or scale of altruism. Similarly, the resource competition experienced at position ***y*** is defined as

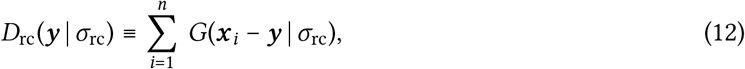

where *σ*_rc_ is the range or scale of resource competition. Note that *D*_rc_(***y*** | *σ*_rc_) is a kernel density estimate based on a Gaussian kernel with band width *σ*_rc_; in the derivation of the LSD the local population density was defined in the same way.

In the simulations, space is discretised using a regular square lattice with lattice constant *δx*, and time proceeds in discrete steps *δt*. The simulation construct the state of the system at time *t* + *δt* through the following sequence of events:

##### 1. Reproduction

Each individual *i* that exists at time *t* reproduces with probability *g*_*i*_*δt*, where the reproduction rate *g*_*i*_ is given by

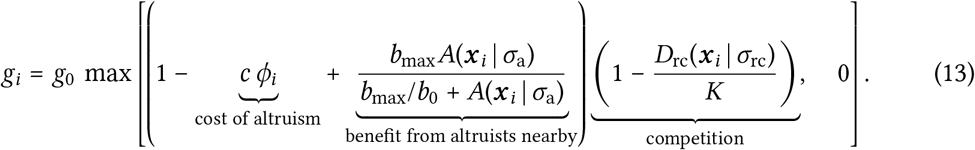

Here, *g*_*0*_ is the basal reproduction rate, *c* the reproductive deficit per investment in altruism, ***x***_*i*_ the position of the individual, and *K* is a factor scaling the local carrying capacity. The benefit of altruism is an increasing, saturating function of the local availability of “public good”: it is 0 if the local environment does not contain any altruists, and *b*_max_ if *A*(***x***_*i*_ | *σ*_a_) → *∞*. When an individual reproduces, a new organism is born and placed at the same lattice point as its parent.

##### 2. Heredity and mutation

The phenotype of the child is copied from its parent, but mutated with probability *µ*. The effect size of the mutation is drawn from an exponential probability distribution with mean *m*; its sign is chosen positive or negative with equal probability. If a mutation would result in a negative phenotype, the phenotype is set to zero.

##### 3. Death

Individuals die with probability *dδt*. The death rate *d* is independent of phenotype.

##### 4. Motility

All individuals are displaced by adding a random number to both coordinates of their position. These random numbers are independently drawn from a discretised normal distribution with standard deviation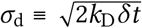. This approximates movement by diffusion, with diffusion constant *k*_D_.

#### Units and parameters

In the formulation above, we are free to choose the scaling of the phenotype *ϕ* this allows one to eliminate one parameter. We choose to set the cost to *c = 1*. We can eliminate another parameter by choosing suitable units of length; we choose to set *σ*_a_ *=* 1, so that all lengths become expressed in units of the interaction range of altruism. A last parameter can be eliminated by choosing convenient units of time; we set the death rate *d =* 1, so that time is measured in generations.

Other parameter values used in the simulations shown in the main text are listed in Table 1. While some parameters were chosen arbitrarily, the following rationale was followed: To resolve the altruistic interactions, the spacing of the lattice needs to be smaller than the range of altruism, *σ*_a_ *=* 1. We therefore set the lattice spacing to *δx* = 0.1. Similarly, to ensure that *g*_*i*_ and *d* can indeed be interpreted as rates, the time step should be chosen such that *g*_*i*_*δt* and *dδt* are considerably smaller than 1. We therefore set *δt =* 0.08. The Turing-like instability causing the colony formation occurs only if the range of competition is significantly larger than the range of altruism (further explored in [29]); we chose *σ*_rc_ = 4. Since the evolution of altruism is favoured by positive genetic assortment (*i.e*., close proximity of parents and offspring) [10], a low diffusion constant was chosen. Lastly, the parameter *b*_0_ was chosen to ensure that the direct benefit reaped by an isolated individual from its own altruism is considerably smaller than the cost incurred. (Otherwise, the term altruism does not apply.) The benefit term in Eq 13 is bounded by *b*_*0*_*A*(***x***_*i*_|*σ*_a_) *= b*_0_*ϕ*_*i*_*/*(*2π*); hence, by choosing *b*_0_ *=* 1, the benefit is guarantied to be at least a factor *2π* smaller than the cost *cϕ*_*i*_ *= ϕ*_*i*_.

**Table 1.**
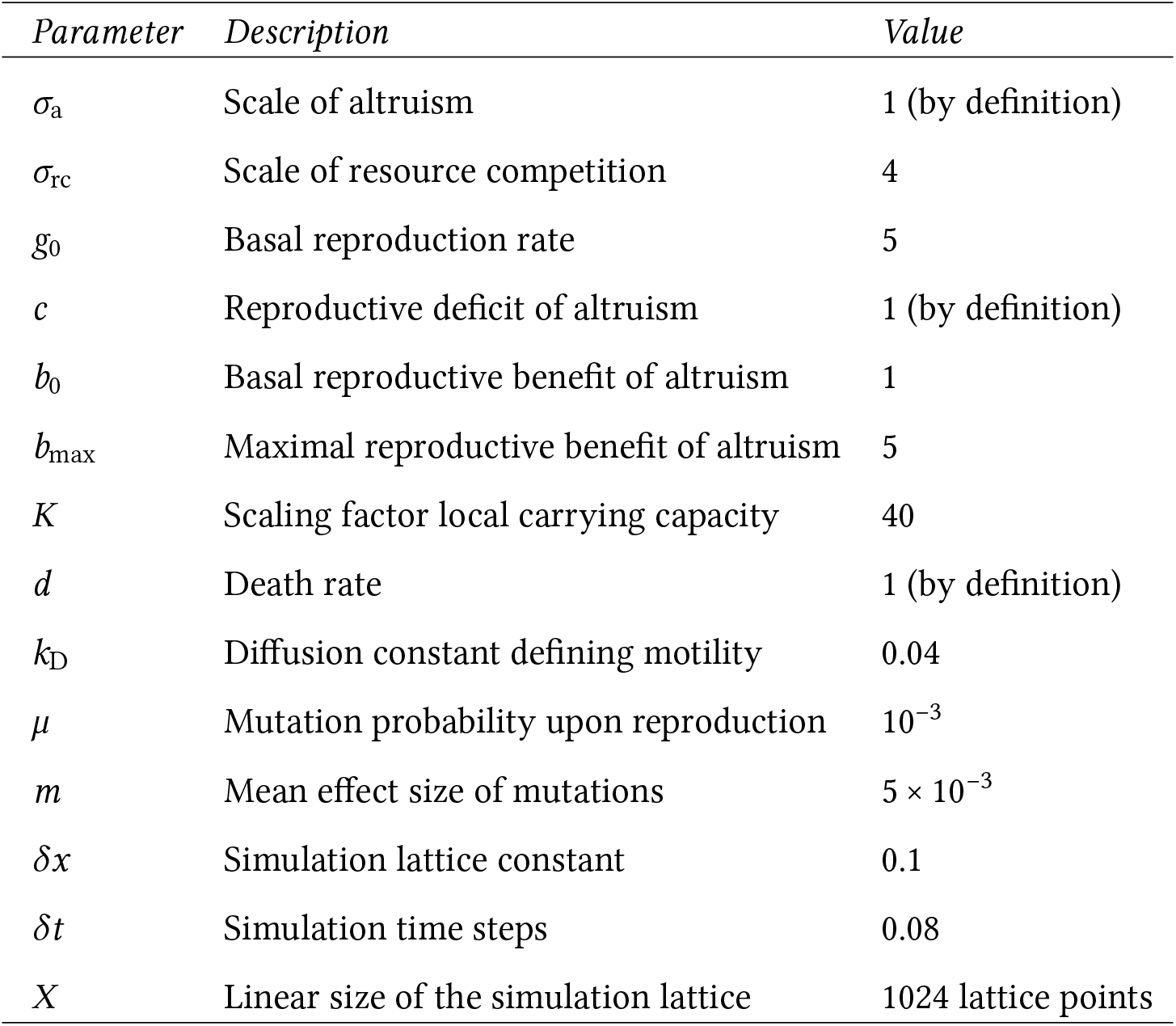
Parameters of altruism model.

### SI-model of the evolution of pathogen transmissibility

As a second example, we consider the evolution of pathogen transmissibility in a spatial SI-model. This model is based on several previous simulation models exploring pathogen evolution, see [32–35].

#### Model description

Consider a population of individuals living on a 2D square lattice. Each lattice point contains at most one individual. Individuals reproduce, die and move randomly. Individuals can be either susceptible to infection with a pathogen, or infected. Infected individuals no longer reproduce, and die at a higher rate than susceptible individuals. The infection spreads locally; the probability that an infected individual transmits the pathogen to a susceptible neighbour depends on the transmissibility of the pathogen that they carry, *ϕ*. We consider the evolution of this transmissibility.

Time in the model progresses in discrete time steps. Every time step, the following series of events takes place:

##### 1. Reproduction

Since each lattice point can be occupied by at most one individual, individuals can reproduce only if one or more of their neighbouring lattice points is empty. Empty lattice points are repopulated by a susceptible individual with probability *γ n*_*S*_, where *γ* is the reproduction rate per susceptible individual and *n*_*S*_ is the number of susceptible individuals among the eight neighbours (including diagonals) surrounding the empty lattice point.

##### 2. Infection

For each susceptible individual *i*, let *I*_*i*_ be the set of infected individuals found on any of the eight lattice points surrounding the susceptible individual’s position. Then individual *i* becomes infected with probability

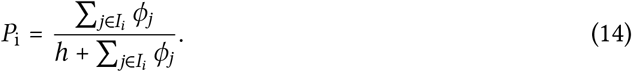

This infection probability is zero when the susceptible individual has no infected neighbours, and approaches 1 if many of the neighbours are infected with highly transmissible pathogens. If an infection takes place, one of the infected neighbours is chosen as the source of this new infection; the probability that a neighbour is chosen is proportional to the transmissibility of its pathogen, *ϕ*_*j*_. The newly infected individual inherits the transmissibility value of the chosen infected neighbour;

##### 3. Mutation

Since we consider mutation of the pathogen rather than of the host individuals, mutations are not coupled to reproduction events of the hosts. Instead, each time step the transmissibility of the pathogen within each infected individual is mutated with a small probability *µ*. This resembles a within-host mutation of the pathogen that (instantly) sweeps the within-host pathogen population. If a mutation occurs, a new transmissibility 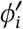 is chosen from a uniform distribution *[ϕ*_*i*_ *− λ; ϕ*_*i*_ + *λ]*. If 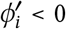, it is set to zero.

##### 4. Death

Susceptible individuals die with probability *δ*_*S*_. Infected individuals die with probability *δ*_*I*_ *> δ*_*S*_.

##### 5. Motility

In a randomly assigned order, the contents of each lattice point are swapped with a randomly chosen neighbour with probability *p*_m_. Individuals can hence change position either because they move themselves (with probability *p*_m_) or because they are chosen in the swapping procedure of one of their neighbouring lattice points (empty or containing an individual). This procedure approximates movement by random walk, while ensuring that the number of individuals does not change and that the rate of movement per individual does not depend on local population density.

#### Parameters

An overview of all model parameters and their values is provided in Table 2. Reproduction and death probabilities were set to values significantly smaller than 1 per time step, so that these probabilities can be interpreted to good approximation as rates of a Poisson process. To allow spatial patterns to arise, a small value was chosen for the motility probability *p*_m_.

**Table 2.** Parameters of SI model.

| *Parameter* | *Description* | *Value* |
| --- | --- | --- |
| $\gamma$ | Reproduction rate (per susceptible per time step) | 0.05 |
| $\delta_S$ | Death rate of susceptible individuals (per time step) | 0.05 |
| $\delta_I$ | Death rate of infected individuals (per time step) | 0.2 |
| $h$ | Scaling factor of infection probability | 20 |
| $p_m$ | Motility probability (per time step) | 0.05 |
| $\mu$ | Mutation rate (per time step) | 0.005 |
| $\lambda$ | Maximal mutation step size | 0.5 |
| | Simulation lattice size | $1024 \times 1024$ lattice points |

#### Scale of emergent spatial patterns

To measure the size of the emergent spatial patterns, we determine the pair correlation function *g*(*r*) of a snapshot of the simulation. This function measures the mean density of individuals at distance *r* from a lattice point, given that this lattice point is occupied by some individual. It thus indicates whether the occupancy of a lattice point correlates with the occupancy of lattice points a distance *r* away. The function is normalised by the average density of the field. Hence, if *g*(*r*) *=* 1, the probability of finding other individuals at distance *r* from a given focal individual is equal to the probability of finding individuals in any random position; if *g*(*r*) > 1 the probability of finding other individuals at distance *r* from a focal individual is larger than one would expect from random sampling of the population.

For patch structures such as the emergent patterns studied here, *g*(*r*) can be approximated by an exponential function

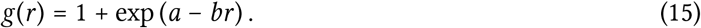

Eq 15 was used to fit the observed pair correlation functions using least-squares fitting. The fitted exponent *b*^−1^ was used as a measure of the size of the patches emerging in the field.

### Implementation of the calculations of *S*_local_(*r*) and *S*_interlocal_(*r*)

In the Methods section, we gave a general analytical derivation of the spatial decomposition of the selection differential *S* into local and interlocal components *S*_local_ and *S*_interlocal_. Below, we describe how these quantities were calculated in practice.

When deriving the spatial decomposition of selection (Methods), we considered a population in continuous space. In the simulations, however, space is discretised. This is easily dealt with by replacing the integral over space in Eq 8 with a double summation over the coordinates of the 2D simulation space.

For both models, simulations were performed using periodic boundary conditions (a common choice for these kinds of simulations). To calculate local population densities (Eq 4) one then must not only take the individuals in the simulation lattice into account, but also their periodic images. The circular convolution theorem provides an efficient way to do this using Discrete Fourier Transforms (DFTs). This is a standard technique from signal processing; we shortly discuss it here for completeness.

For ease of notation we here present derivations for a 1D space; the 2D case follows analogously.

Assume that a sequence o_0_, o_1_, …, o_*X−*1_ specifies the occupancy (number of individuals) at discrete positions *j* ∈ {0, 1, …, *X −* 1} in a simulation field, and let 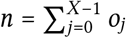 be the total number of individuals. Because we assumed periodic boundary conditions, we should consider this finite field to represent a finite stretch taken from an infinite field that is periodic with period *X*; that is, the full field, including periodic images, is a sequence *õ*_*j*_ with *j* ∈ ℤ defined by *õ*_*j*_ *= o*_(*j* mod *X*)_.

On the discretised space, the kernel function is represented by an infinite non-negative sequence *k*_*j*_ with *j* ∈ ℤ, whose sum converges to 1. Because the kernel function is a function of distance, it is symmetric, *i.e*., *k*_*−j*_ *= k*_*j*_. The kernel density *d*_*j*_ for positions *j* ∈ {0, 1, …, *X* − 1} (*c.f*. Eq 4) is now given by

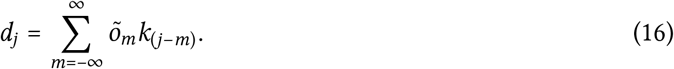

Eq 16 can be rewritten as follows:

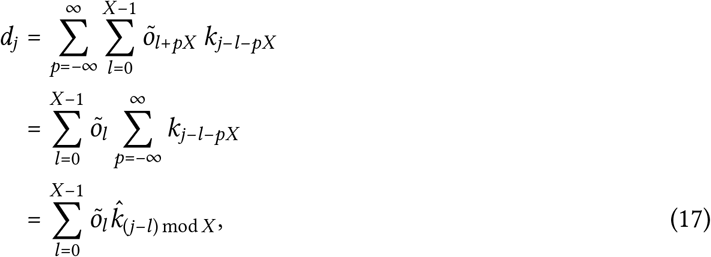

where we introduced the *periodic summation* 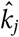 for *j* ∈ {0, 1, …, *X* − 1} as

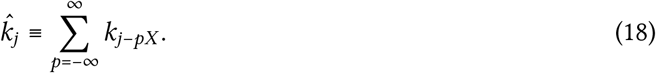

The sequences *õ*_*j*_ and 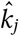 are both periodic with period *X*. Eq 17 is the *circular convolution* of these two sequences, which we denote by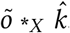. By the *circular convolution theorem*, the DFT of 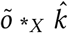 is equal to the element-wise product of the DFTs of *õ* and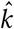, *i.e*.,

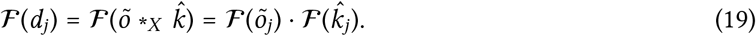

Using Eq 18 and 19, we can calculate the local density *d*_*j*_ by

1. calculating the periodic summation 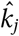 of *k*_*j*_;
2. calculating the DFTs of 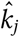 and *o*_*j*_, and calculating their element-wise product;
3. calculating the inverse DFT of this element-wise product, which yields *d*_*j*_.

Because algorithms for Fourier transformation are highly efficient [55], this procedure allows us to rapidly solve local densities for varying scales *r*. Fourier transformations were performed using the fftw3-library [55].

The local mean of a property *c* of individuals, {*c* | *j, r*}_l_ (Eq 6), can be calculated in a very similar way. Let 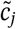 be the sequence of the sum of *c*-values of individuals at position *j*. In terms of this sequence, we can write

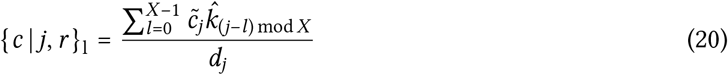

In other words, the total local value of *c*, {*c* | *j, r*}_l,tot_ ≡ *d*_*j*_ {*c* | *j, r*}_l_, is given by the circular convolution of the sequences 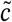 an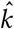, and can hence be easily calculated using Fourier transformation as described above. To find the local mean, this total local value is divided by the local density, *d*_*j*_. Note that, for some positions *j*, the local density might be very small or zero. To avoid numerical problems, division by local density is treated with caution in the calculations of *S*_local_(*r*) and *S*_interlocal_(*r*) (see below).

By definition, *S*_local_(*r*) is the spatial mean of the LSDs, weighted by the local mean fitness {*w*}_l_. It is therefore possible to calculate *S*_local_(*r*) directly from the local selection differential, *S*_1_((*x, y*) | *r*):

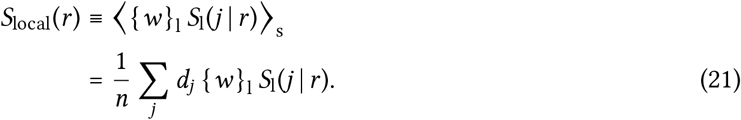

However, noting that 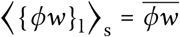 we can also calculate *S*_local_(*r*) using Eq 10:

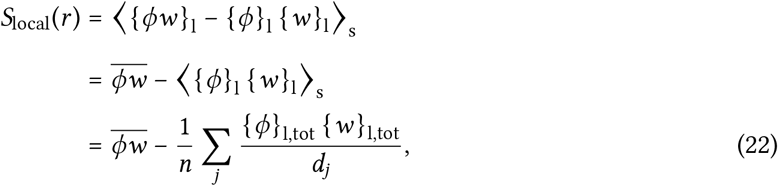

which is less likely to cause numerical inaccuracies (if the local density *d*_*j*_ *=* 0 for some position *j*, the corresponding term in the summation is set to zero).

Using the definition of Eq 9, the value of *S*_interlocal_(*r*) can be calculated directly as the covariance between {*ϕ*| *j, r*}_l_ and {*w* | *j, r*}_l_, in which each position *j* is weighted by the local density *d*_*j*_. However, we can again derive an expression that is numerically more stable:

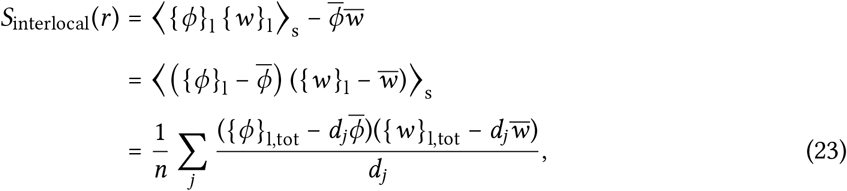

where again the summation term for index *j* is set to zero if the local density *d*_*j*_ *= 0*. Eq 22 and 23 were used in the calculations.

